# Metabolic engineering of *Escherichia coli* strains for the *in vivo* synthesis of GP-mediated oligosaccharides

**DOI:** 10.64898/2026.02.23.707450

**Authors:** Pietro Tedesco, Julien Durand, Laurence Tarquis, Gabrielle Potocki-Veronese, Fabien Létisse

**Affiliations:** Institut de Pharmacologie et de Biologie Structurale (IPBS), Université de Toulouse, CNRS, UPS, Toulouse, France; TBI, Université de Toulouse, CNRS, INRAE, INSA, 31077 Toulouse, France; Stazione Zoologica Anton Dohrn, Department of Ecosustainable Marine Biotechnology, Naples, Italy; Sweetech SAS, INSA, TBI, 135 Av. de Rangueil, 31077 Toulouse, France

**Keywords:** Reverse phosphorolysis, Glycoside phosphorylases, β-mannobiose, Metabolic engineering, Carbon flux control, Growth–production decoupling

## Abstract

Enzymatic synthesis of rare disaccharides by reverse-phosphorolysis is a potentially sustainable route to produce high-value glycosides for human health and nutrition. We report the metabolic engineering of *Escherichia coli* for *in vivo* production of β-mannobiose with different osidic linkages from hexose sugars. We demonstrate production of β-1,2-mannobiose with this approach as proof of concept. Phosphotransferase system (PTS) inactivation enables import of non-phosphorylated mannose *via* heterologous permease GalP, restoring growth on mannose in a PTS⁻ background and allowing mannose into the reverse biosynthetic pathway. Deletion of *pfkA*, which promotes intracellular accumulation of key sugar phosphates (G1P, M1P), establishes a favorable metabolic chassis for oligosaccharide production using glycoside-phosphorylases.

Using this chassis, we expressed two β-mannoside phosphorylases to enable the direct production of β-1,2- and β-1,4-mannobiose from mannose. The same chassis was also employed for laminaribiose production through the expression of a laminaribiose-phosphorylase. *pfkA* deletion significantly increased product titer (> 0.6 g·L^−1^) and yield (up to 9% g/g mannose), highlighting a favorable redistribution of carbon fluxes toward disaccharide formation. Moreover, a combination of mixed-substrate cultures using glycerol as carbon and energy source and further metabolic engineering enabled partial growth-production decoupling, redirecting mannose utilization primarily toward product synthesis, with a yield of 60%. These results demonstrate the modularity and efficiency of the proposed platform for fermentative production of non-conventional oligosaccharides and expand the scope of metabolic engineering strategies for glycoside biosynthesis.

## 1. Introduction

Oligosaccharides are a highly diverse class of biomolecules consisting of linear or ramified chains of glycosyl moieties or glycosyl derivatives. Oligosaccharides play a multitude of biological roles in cellular processes, the modulation of immune response and host-microbe interactions (Varki, 2017), and have thus found a broad range of applications in the food (functional foods, prebiotics or as additives) and health (as drugs, drug carriers or vaccine intermediates) industries (Guan et al., 2019; Mensink et al., 2015; Swanson et al., 2020; Zhang et al., 2019, 2018; Zheng et al., 2020), with ongoing scientific and economic interest in improving their production. Oligosaccharides can be produced enzymatically and chemo-enzymatically by exploiting the natural diversity and selectivity of enzymes to enlarge the panel of glycan structures that can be produced, avoiding multiple chemical synthesis steps (Báti et al., 2019; Li et al., 2019; Reetz, 2013; Rudroff et al., 2018). Four main types of enzymes have been used to efficiently catalyze oligosaccharide production: glycoside-hydrolases (GHs) and glyco-synthases, transglycosylases (TGs), glycosyltransferases (GTs) and glycoside-phosphorylases (GPs) (Faijes et al., 2019). GPs are reversible enzymes that catalyze both the breakdown of osidic linkages in the presence of phosphate (in what is commonly referred to as the reverse-phosphorolysis reaction) and the synthesis of glycosides from a glycosyl-phosphate and an acceptor (the reverse phosphorolysis reaction) (Puchart, 2015). However, regardless of the classes and mechanisms involved, use of glycoside synthesizing enzymes in *in vitro* processes is limited by enzyme production and protein purification costs, loss of enzyme activity (in storage and under reaction conditions), and also in most cases, by substrate costs. Specifically, glycosyltransferase-based approaches require expensive nucleotide-sugars and glycosynthase-based methods require chemically modified glycosyl donors that are difficult to produce with high yields at a reasonable cost (Cobucci-Ponzano and Moracci, 2012). *In vivo* synthesis using microorganisms that express recombinant glycoside synthesizing enzymes has therefore attracted attention for its potential to circumvent these limitations. This approach is based on the ability of the engineered cells to produce the precursor substrates (activated sugars) from affordable carbon sources as well as the catalysts and the targeted products, leading to a more economically sustainable process. Microbial production has proved particularly popular in recent years for human milk oligosaccharide production, with GTs as biocatalysts (Lu et al., 2021; Meng et al., 2023). The use of other catalysts has been far more limited however; for instance, GPs have only been used *in vivo* alongside GTs (De Bruyn et al., 2015), in the phosphorolysis direction, to increase the pool of sugar phosphates through the phosphorolysis of low-cost oligosaccharides, the sugar phosphates being themselves transformed into nucleotide sugars, the substrates of GTs. Nonetheless, *in vivo* reverse-phosphorolysis has been reported in *Mycobacterium smegmatis* strains, which can use native GPs for the synthesis of energy storage glycosides such as glycogen from sugar phosphates (Elbein et al., 2010). Recently, Tian *et al*. (Tian et al., 2020) reported the production of manno-oligosaccharides in a mannose fermentation system using an engineered strain of *Corynebacterium glutamicum*.

To date however, oligosaccharide synthesis through GP-based reverse-phosphorolysis has yet to have been achieved with *E. coli*, the preeminent host for oligosaccharide production using GTs (Bych et al., 2019). One explanation for this is the low intracellular availability of the substrates in *E. coli*. For oligosaccharide synthesis indeed, GPs require sugar phosphates as donors and non-phosphorylated sugars as acceptors, and while the former are abundant in *E. coli* cells, non-phosphorylated monosaccharides are lacking in native strains because most carbohydrates are internalized by the phosphoenolpyruvate–carbohydrate phosphotransferase system (PTS) (Escalante et al., 2012), preventing the use of GPs in the reverse-phosphorolysis direction.

In this study, we tackled the problem of limited precursor availability in *E. coli* using a metabolic engineering strategy focused on oligosaccharide production *via* GPs. Central to this strategy is the decoupling of sugar import from phosphorylation through the replacement of the native PTS (which internalizes the carbon source and monosaccharide acceptors) by a permease-based transport mechanism. Further deletions in central metabolism lead to the accumulation of the sugar phosphate intermediates required to produce oligosaccharides, which are then easily recovered from the culture medium. Further engineering allowed the monosaccharide acceptor flux to be channeled almost completely toward oligosaccharide synthesis, using glycerol as carbon and energy source. As proof of concept of this approach, we report the production of various oligosaccharides of interest for nutrition and health, in particular β-1,2-mannobiose, an antigenic motif of the pathogenic yeast *Candida albicans* which is very difficult to chemically synthesize cost-effectively at large-scale (Rahkila et al., 2014), hindering the development of candidiasis vaccines.

## 2. Material and methods

### 2.1 Chemicals

β-1,4-mannobiose and laminaribiose were purchased from Biosynth (Bratislava, Slovakia). β-1,2-mannobiose was purchased from Omicron Biochemicals (South Bend, IN, USA). LC-MS grade solvents (methanol, acetonitrile) were obtained from Instrumentation Consommables et Service (ICS, Lapeyrousse-Fossat, France).

### 2.2 Strain construction

Strain MDO, constructed from strain ZLKA, was provided by Dr Éric Samain (CERMAV, Grenoble, France). Strain ZLKA is a derivative of *Escherichia coli* K12 strain DH1 (*endA1 recA1 gyrA96 thi-1 glnV44 relA1 hsdR17*), constructed by disrupting genes *lacZ*, *nanKETA*, and *lacA* as previously described (Fierfort and Samain, 2008). Strain MDO involves three additional mutations (*melA, wcaJ, mdoH*) and insertion of a Plac promoter upstream of *gmd* (encoding for GDP-mannose 4,6-dehydratase) (Samain laboratory, unpublished research). As a result, the *manB* gene for phosphomannomutase, which is located downstream from the *gmd* gene, is inducible by ITPG. The complete genotype of the parental MDO strain is: *endA1 recA1 gyrA96 thi-1 glnV44 relA1 hsdR17 lacZ-wcaF::Plac nanKETA lacA melA wcaJ mdoH*.

Selected genes (*ptsG*, *manXYZ*, and *pfkA*) were deleted using the PKO3 gene knock-out protocol (Link et al., 1997). PKO3 plasmid was obtained from the Church laboratory (Harvard Medical School). Briefly, flanking regions upstream and downstream of genes selected for deletion were amplified by PCR and cloned together into the *BamHI SalI* sites of the suicide vector pKO3. For *ptsG*, the upstream fragment was cloned with primers P1 and P2 and the downstream fragment with primers P3 and P4 (see Supplementary Table 1), resulting in the deletion of a 0.351 kb DNA segment between nucleotides 568 and 919 of *ptsG*. For *manXYZ*, the upstream fragment was cloned with primers P5 and P6 and the downstream fragment with primers P7 and P8, resulting in the deletion of a 1.219 kb segment between nucleotides 332 of *manX* and 522 of *manY*. For *pfkA*, the upstream fragment was cloned with primers P9 and P10 and the downstream fragment with primers P11 and P12, resulting in deletion of a 0.396 kb DNA segment between nucleotides 146 and 542 of *pfkA*.

The genes *maa* and *manA* were deleted using the Datsenko and Wanner protocol (Datsenko and Wanner, 2000). Briefly, a DNA cassette containing the gene for kanamycin resistance and flanked by Flippase Recognition Target (FRT) sites was amplified using the plasmid pKD4 as template, with 50 bp of the 5’ and the 3’ ends of the gene at the extremities. For *maa* and *manA,* the primer couples were P13/P14 and P15/P16, respectively.

The resulting strains were transformed with plasmid pKD46 at 30°C on LB agar plates supplemented with ampicillin at 50 µg/mL. Transformants carrying pKD46 plasmid were grown in 5-ml LB cultures with ampicillin and L-arabinose (0.2% w/v) at 30°C to an optical density at 600 nm (OD_600_) of 0.6 and then made electrocompetent by concentrating 100-fold and washing three times with ice-cold 10% glycerol. Electroporation was done using a Cell-Porator device with a voltage booster and 0.1 cm chambers according to manufacturer instructions, with 50 µL of cells and 200 ng of PCR product. Shocked cells were added to 1 ml of SOC medium, incubated for 2 h at 37°C, and then spread onto agar to select Km transformants. Once colonies lacking the gene had been identified, the cassette was removed by transforming one of the positive colonies with the plasmid pCP20. This plasmid is an ampicillin resistance plasmid with temperature-sensitive replication and thermal induction of FLP synthesis. KmR mutants were transformed with pCP20, and ampicillin-resistant transformants were selected at 30°C, after which a few were colony-purified once, non-selectively, at 43°C and then tested for loss of all antibiotic resistances.

All PCR reactions were performed with NEB phusion DNA polymerase according to manufacturer instructions, adapting cycles to primers’ melting temperatures and fragment lengths.

### 2.3 Plasmid construction

#### 2.3.1 Cloning of monosaccharide transporter

A 1.471 kb DNA fragment containing the sequence of the *galP* gene was amplified by PCR using the genomic DNA of Escherichia coli K12 as a template and primers P17 and P18. The amplified fragment was then cloned into the *SalI* and *XbaI* sites of the pWKS130 expression vector to form pWKS-GalP {Citation}.

#### 2.3.2 Cloning of GPs

The genes encoding the Teth514-1788 protein from *Thermoanaerobacter* sp. strain X-514 (Teth514-1788, GenBank accession number ABY93073.1 (Ahmadipour et al., 2023)), a β-1,2-mannoside-phosphorylase belonging to the GH130 family in the CAZy classification (http://www.cazy.org/) (Lombard et al., 2014), and the gene encoding the ACL0729 protein, a laminaribiose-phosphorylase from *Acholeplasma laidlawii* strain PG-8A (ACL0729, GenBank accession number ABX81345.1 (Nihira et al., 2012)), belonging to the GH94 family in the CAZy classification (http://www.cazy.org/) (Lombard et al., 2014), were synthesized and cloned into the pBAD HisA vector by GeneCust (GeneCust, Boynes, France) between the NcoI and XhoI restriction sites.

The gene encoding the Uhgb_MP β-1,4-mannoside-phosphorylase from an uncultured human gut bacterium, belonging to the GH130 family (Uhgb_MP, GenBank accession number ADD61463.1 (Ladevèze et al., 2013)), was cloned into the pBAD HisA vector using the primers P19 and P20.

Cloning was performed according to manufacturer’s instructions (Invitrogen Corp., Carlsbad, CA, USA). All plasmid constructs were verified by sequencing, and protein expression profiles were assessed by SDS-PAGE electrophoresis using Any kD gels from Bio-Rad as follows. Cell pellets were lysed in 5 mM Tris-HCl buffer containing lysozyme (Euromedex, ref. 5934-C) at 0.5 mg·mL⁻¹ and DNase (NEB, ref. M0303L) at 50 µL (100 U) per 100 mL of 5 mM Tris-HCl buffer. All pellets were resuspended in buffer to an OD_600_ of 45, incubated for 30 min at 37 °C, and then frozen at –20 °C. Samples were thawed and centrifuged for 25 min at 5000g. Fifteen microliters of each supernatant were mixed with 5 µL of blue loading dye (NEB, ref. 50994905), incubated for 10 min at 95 °C, and then loaded onto the gel. Electrophoresis was carried out for 35 min at 100 V in an electrophoresis tank to confirm that the proteins were indeed produced in the transformed PFKA1 strain and had the expected molecular masses.

### 2.4 Media and growth conditions

*Escherichia coli* strains C for DNA manipulation and strain construction were grown at 37°C or 42° or 30°C in LB media (10 g·L^−1^ tryptone, 5 g·L^−1^ yeast extract, 10 g·L^−1^ NaCl) as indicated. All other growth strains were grown in minimal medium M9. M9 salts: Na_2_HPO_4_ 12H_2_O 17.4 g·L^−1^, KH_2_PO_4_ 3.02 g·L^−1^, NaCl 0.51 g·L^−1^, NH_4_Cl 2.04 g·L^−1^. Trace metals salts: Na_2_EDTA 2 H_2_O 15 mg·L^−1^, ZnSO_4_ 7 H_2_O 4.5 mg·L^−1^, CoCl_2_ 6H_2_O 0.3 mg·L^−1^, MnCl_2_ 4H_2_O 1 mg·L^−1^, H_3_BO_3_ 1 mg·L^−1^, Na_2_MoO_4_ 2 H_2_O 0.4 mg·L^−1^, FeSO_4_ 7 H_2_O 3 mg·L^−1^, CuSO_4_ 5 H_2_O 0.3 mg·L^−1^. MgSO_4_ 0.5 g·L^−1^ CaCl_2_ 4.38 mg·L^−1^, Thiamine hypochloride 0.1 g·L^−1^.

*E. coli* strains were grown in M9 minimal medium complemented with the desired carbon sources. Cultures were performed in duplicate or triplicate in shake-flasks at 37 °C and 200 rpm, using a working volume of 250 mL containing 3 g·L⁻¹ of selected carbon source. Growth was monitored by measuring the OD_600_ using a Genesys 6 spectrophotometer (Thermo, USA). Kanamycin and ampicillin were added at a final concentration of 50 µg·mL^−1^ to ensure propagation of plasmids. IPTG (final concentration 60 µM) and arabinose (final concentration 1 mM) were added to the media to induce protein expression. For bioreactor experiments, the concentrations of M9 salts were changed to: KH_2_PO_4_, 3.02 g·L^−1^; NaCl, 0.51 g·L^−1^; NH_4_Cl, 2.04 g·L^−1^; (NH_4_)_2_SO_4_, 5 g·L^−1^. All the other components were as described above. Bioreactor cultures were performed in 500 ml of M9 medium complemented with desired carbon source. Parameters were set and monitored using a Multifors bioreactor system (Infors,Switzerland). The percentages of CO2 and O2 in the gas output were determined using a Dycor ProLine Process Mass Spectrometer (Ametek, DE, USA).

### 2.5 Metabolomics analysis

#### Sampling and metabolite extraction

Samples (100 µL) of an overnight culture of liquid LB with the desired strains were used to inoculate 50 mL baffled shake flasks containing 10 mL of M9 medium supplemented with 3 g·L^−1^ of glucose. The flasks were incubated for 24 h at 37°C with orbital shaking at 220 rpm. Cells were harvested by centrifugation (Sigma, Seelze, Germany) for 10 min at 2000g at room temperature, washed with five times diluted M9 and used to inoculate 50 mL baffled shake flasks at OD_600_ = 0.1 containing 50 mL of diluted M9 supplemented with 3 g·L^−1^ of glucose and incubated at 37 °C at 220 rpm. Cell growth was monitored by measuring the OD_600_ with a cell density meter (Ultrospect 10, Amersham BioSciences). Sampling was performed using the fast filtration method [25]. When cultures reached the exponential phase, (OD_600_ of 1–1.5) 120 μL of broth was harvested by vacuum filtration (Sartolon polyamide 0.2 lm). Cells were washed with 1 ml of medium with five times reduced concentrations of phosphate and sulfate salts and of carbon source. All the other components of the medium were unchanged in the washing solution. The filters were rapidly placed in liquid nitrogen and stored at −80°C until further metabolite extraction. For metabolite extraction, the filters were incubated in closed glass tubes containing 5 ml of a precooled acetonitrile/methanol/H2O (4:4:2) solution and 60 µL of fully 13C-labeled cell extract were added to each tube as internal standard. The tubes were incubated for 1 h at −20 °C, the filters were removed, and the supernatants were evaporated under vacuum (SC110A SpeedVac Plus, ThermoSavant, Waltham, MA, USA) for 12 h and then stored at −80 °C until further treatment.

### IC-ESI-HRMS analysis of central metabolites

Samples were dissolved in 120 µL of mQ water and then centrifuged at 10000g at 4°C for 10 min to remove cell debris. The supernatant was then analyzed by ion chromatography (Thermo Scientific Dionex ICS-5000+ system, Dionex, Sunnyvale, CA, USA) mass spectrometry, on LTQ Orbitrap device (Thermo Fisher Scientific, Waltham, MA, USA) equipped with an electrospray ionization probe. The ion chromatography method was adapted from Kiefer et al. (Kiefer et al., 2007). The KOH gradient was modified as follows: 0 min, 0.5 mM; 1 min, 0.5 mM; 9.5 min, 4.1 mM; 12.5 min, 30 mM; 24 min, 50 mM; 36 min, 60 mM; 36.1 min, 90 mM; 43 min, 90 mM; 43.5 min, 0.5 mM; 48 min, 0.5 mM. The gradients were all linear. A Dionex anionic electrolytically regenerated suppressor (300–2 mm) was used for background suppression. The device was operated at a constant electrolysis current of 87 mA, in external-water mode with ultrapure water, and regenerant was delivered using an external AXP pump at a flow rate of 1 mL/min. The volume of the injected sample was 35 μL in all cases. Mass spectrometry analysis was performed in the negative FTMS mode at a resolution of 30,000 in fullscan mode, with the following source parameters: capillary temperature, 350°C; source heater temperature, 300°C; sheath gas flow rate, 50 a.u. (arbitrary units); auxiliary gas flow rate, 5 a.u; S-Lens RF level, 60%; and ion spray voltage, 3.5 kV. The data were acquired using the Xcalibur software and processed using the software TraceFinder 3.1 (Thermo Fisher Scientific, Waltham, MA, USA). 6-phosphogluconate (6PG) fructose-6-phosphate (F6P), fructose-1,6-bisphosphate (FBP), Glucose-6-phosphate (G6P), combined pools of glucose-1-phosphate and mannose-1-phosphate (G1P/M1P), glucosamine-6-phosphate(GlcN-6P), N-Acetylglucosamine-1-phosphate (GlcNAc-1-p), combined pools of xylulose-5-phosphate, ribose-5-phosphate, ribulose-5-phosphate (P5P), phosphoenolpyruvate (PEP) and sedoheptulose-7-phosphate (S7P) were injected as pure standards, and the isotope dilution mass spectrometry (IDMS) method was used to ensure accurate quantification (Wu et al., 2005). The absolute intracellular concentration for each metabolites in the different strains was calculated from these calibration curves, assuming a cell volume of 1.77 ml/gDW (Chassagnole et al., 2002).

### 2.6 NMR analysis

Exocellular metabolites were identified and quantified by nuclear magnetic resonance (NMR). Broth samples were collected at different times and centrifuged. The supernatants (500 μL) corresponding to the culture medium, were mixed with 100 μL of D2O containing 2.35 g·L^−1^ of TSPd4 (3-(trimethylsilyl)-1-propanesulfonic acid-tetra deuterated) as internal compound. Proton NMR spectra were recorded on an Avance 500 MHz NMR spectrometer equipped with a 5 mM BBI probe head (Bruker, Rheinstatten, Germany). Spectra processing and metabolite quantification were performed using Topspin 3.1 (Bruker, Rheinstatten, Germany). The concentrations of extracellular metabolites were determined using TSP as internal standard (Turbitt et al., 2020).

### 2.7 HPAEC-PAD ANALYSIS

Culture supernatants (1 mL) were heated at 95 °C for 10 min, centrifuged for 10 min at 15000g, and the resulting supernatants were passed through a 0.22 µm membrane filter. Filtered samples were diluted 25-fold in ultrapure water prior to chromatographic analysis. Carbohydrate and related compounds were separated on a Dionex CarboPac PA100 column (2 × 250 mm; Thermo Fisher Scientific) using a 0.25 mL·min⁻¹ elution gradient with the following eluents: A, H₂O; B, 150 mM NaOH; and C, 150 mM NaOH supplemented with 500 mM sodium acetate. The gradient program was: 0–3 min, 50% A / 50% B; 3–6 min, 100% B; 6–12 min, 50% B / 50% C; 12–15 min, 5% B / 95% C; followed by re-equilibration at 50% A / 50% B from 15–23 min. Detection was carried out using a Dionex ED40 electrochemical detector equipped with a gold working electrode and an Ag/AgCl reference electrode.

## 3. Results

### 3.1 Metabolic engineering strategy for mannobiose bioproduction

The strategy devised to enable *in vivo* reverse-phosphorolysis by GPs for mannobiose production in *E. coli* is presented in Figure 1. To increase the intracellular availability of unphosphorylated mannose, the native PTS, responsible for coupled uptake and phosphorylation of glucose and mannose, was inactivated by deleting the *ptsG* and *manXYZ* genes. However, since this severely impairs growth on glucose and mannose (Hernández-Montalvo et al., 2003; Plumbridge, 1998), the *galP* gene encoding galactose permease was overexpressed from a plasmid. Importantly, GalP facilitates the import of both mannose (Henderson et al., 1984)and glucose (Hernández-Montalvo et al., 2003) in their unphosphorylated forms, allowing cytosolic accumulation of free hexoses. Mannose is subsequently phosphorylated to mannose-6-phosphate (M6P), presumably by glucokinase (*glk*) (Honghong et al., 2011). From this branch point, M6P can either be converted to fructose-6-phosphate (F6P) by mannose-6-phosphate isomerase (ManA), thereby entering central carbon metabolism, or be converted directly to mannose-1-phosphate (M1P) by phosphomannomutase (ManB), generating the activated donor substrate required for reverse-phosphorolysis. To further favor M1P accumulation, the concentrations of phosphorylated intermediates were altered by deleting the phosphofructokinase gene *pfkA*, thereby modulating intracellular sugar phosphate pools. The assumption underlying this channeling of the carbon flux toward mannobiose biosynthesis is that the disaccharide product is efficiently exported, minimizing feedback inhibition and enabling sustained production.

**Figure 1.**
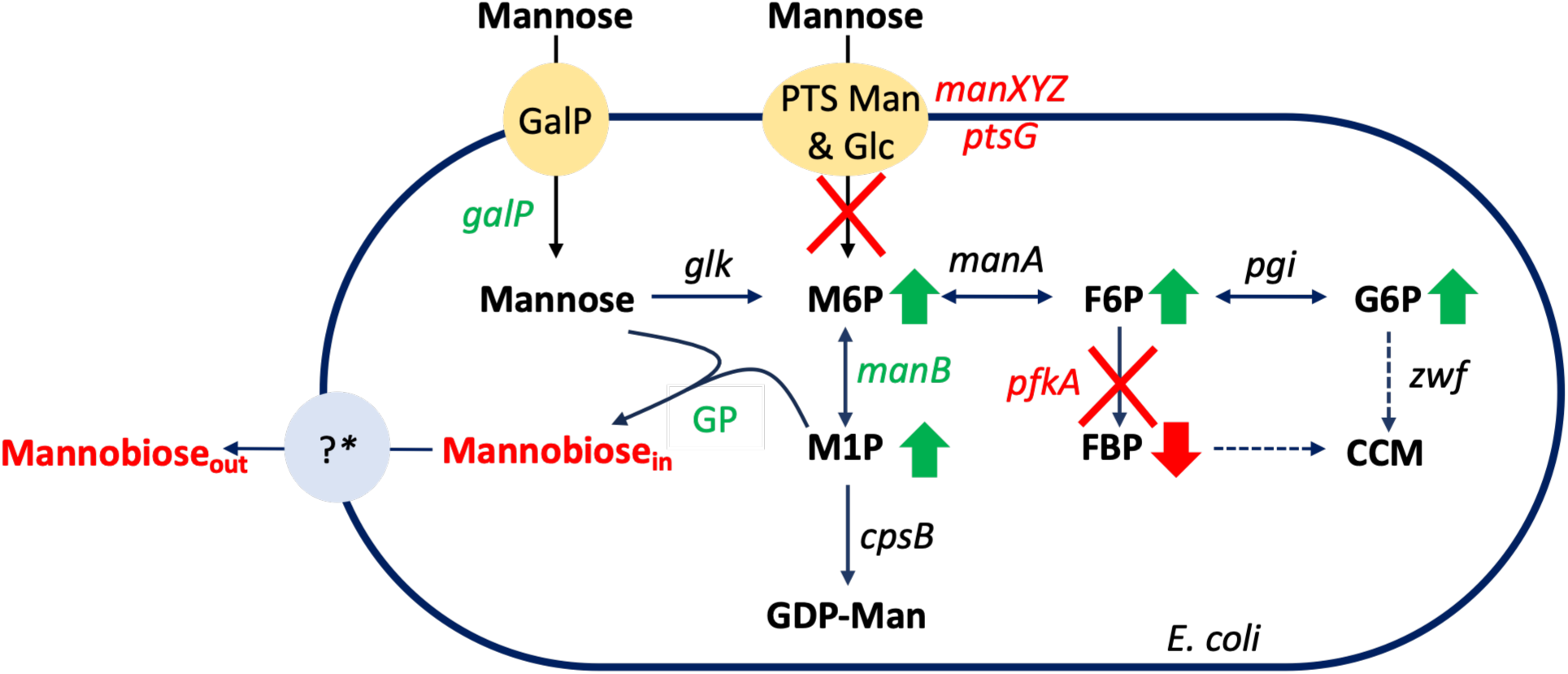
Metabolic engineering of *E. coli* for the synthesis of mannobiose by glycoside-phosphorylases. The strategy involves several gene deletions (indicated in red) and overexpressions (indicated in green). The relative concentrations of intracellular metabolites expected to change upon *pfkA* deletion are indicated by green arrows (for increases) or red arrows (decreases). *Abbrevations: GalP, galactose permease; PTS, phosphotranferase system; manXYZ, mannose specific PTS enzyme II; ptsG, glucose specific PTS enzyme IIBC; glk, glucokinase; manA, phosphomannose isomerase; pgi, phosphoglucose isomerase; manB (cpsG), phosphomannomutase; pfkA, phosphofructokinase I; zwf, glucose-6-phosphate dehydrogenase; cpsB, mannose-1-phosphate guanylyltransferase; GP, glycoside-phosphorylase (mannoside-phosphorylase); M6P, mannose-6-phosphate; F6P, fructose-6-phosphate; G6P, glucose-6-phosphate; M1P, mannose-1-phosphate; FBP, fructose-1,6-biphosphate, GDP-Man, GDP-mannose; CCM, central carbon metabolism.* * *The mannobiose export system is unknown*.

### 3.2 Monosaccharide import through galactose permease in PTS mutant strains

Our approach to mannoside production requires the import of unphosphorylated mannose into the cytoplasm to serve as a substrate (acceptor) for the recombinant mannoside-phosphorylases expressed from plasmids in the chassis strains (Figure 1). We therefore inactivated the PTS, which normally imports and phosphorylates mannose. In *E. coli* however, while the mannose PTS is the primary route for mannose uptake, the glucose PTS can also (less efficiently) transport mannose (Jahreis et al., 2008). We therefore used the MDO strain (Pedersen et al., 2022) (Table 1), based on *E. coli* K-12 strain DH1, previously engineered for oligosaccharide production. This strain has deletions in several galactosidase-encoding genes (*lacZ*, *lacA*, *melA*) and in genes (*nanATEK*, *wcaJ*, *opgH*) involved in the biosynthesis of non-essential oligosaccharides that could interfere with mannoside production. It also has a *Plac* promoter upstream of the *gmd* gene, enabling IPTG-inducible overexpression of downstream genes, including *manB* (*cpsG*), which encodes a phosphomannomutase responsible for the reversible conversion of mannose-6-phosphate (M6P) into mannose-1-phosphate (M1P).

**Table 1:**
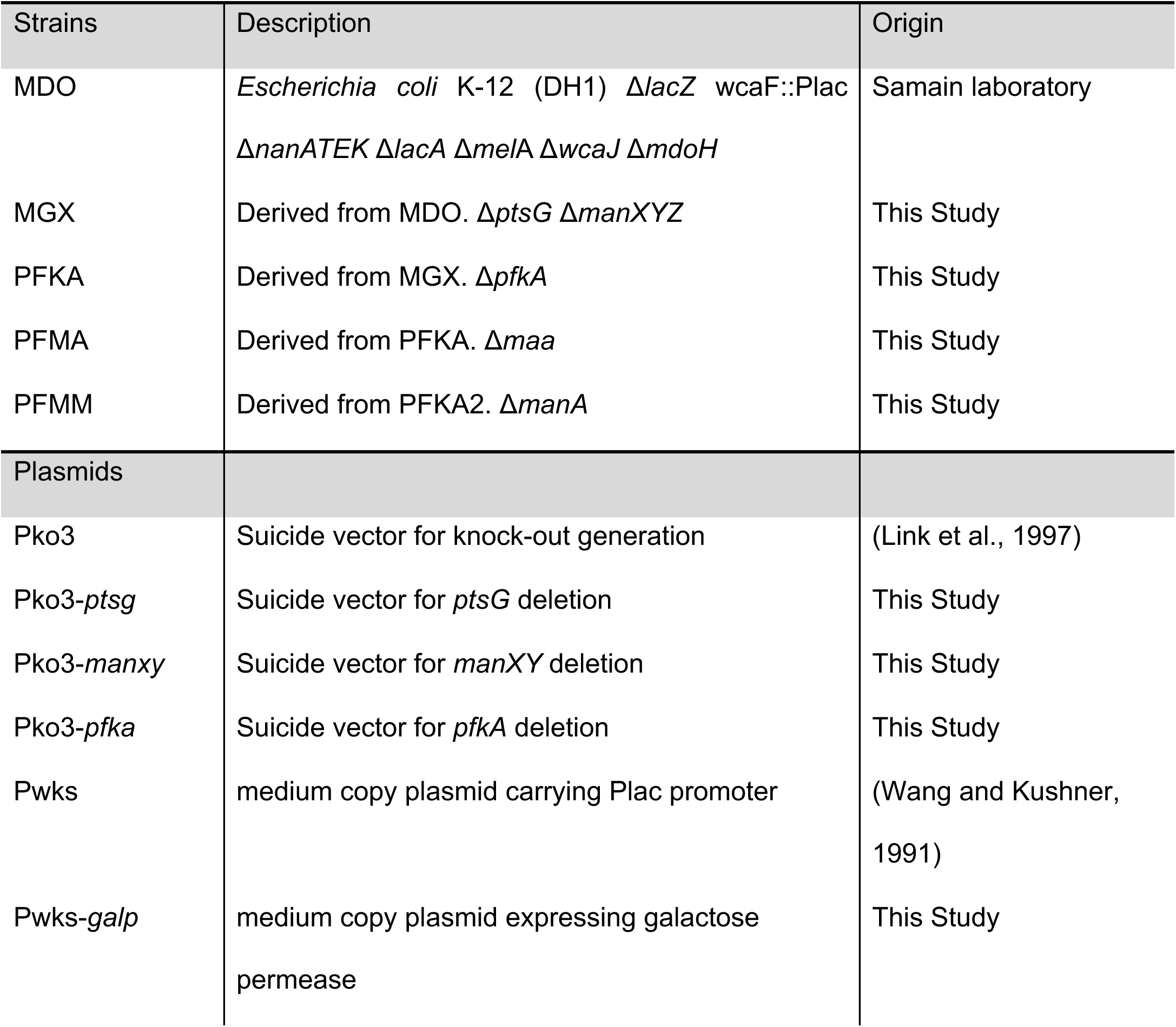
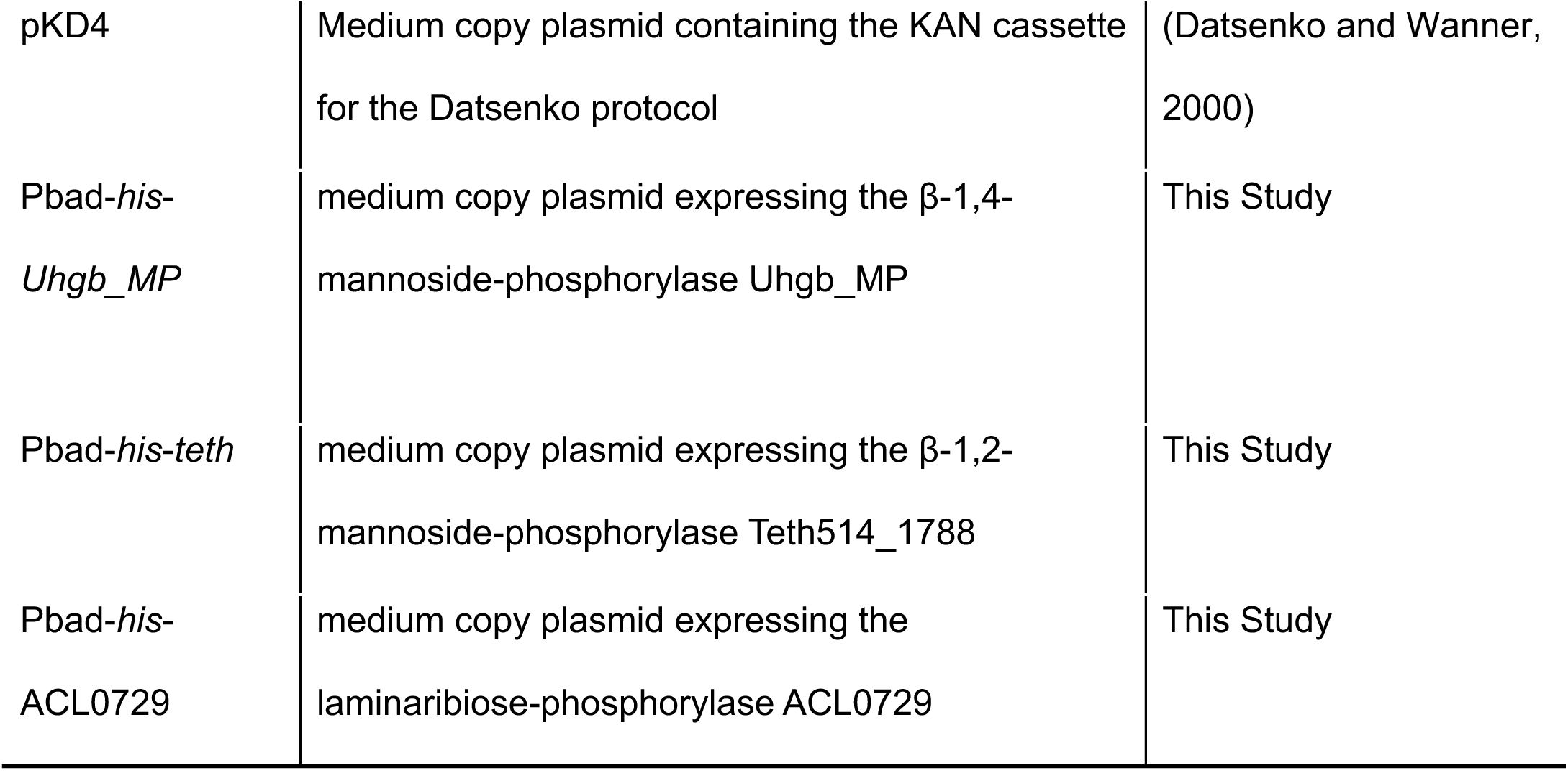
List of strains and plasmids used in this study.

It is well established that PTS transporters are the main routes for glucose and mannose uptake in *E. coli*. These systems mediate the simultaneous import and phosphorylation of sugars, using phosphoenolpyruvate (PEP) as the phosphate donor. To promote the uptake of unphosphorylated mannose, we deleted *ptsG* and *manXYZ*, which respectively encode the IIB subunits of glucose and mannose specific PTS enzyme II, in the MGX strain (Table 1). However, PTS deletion severely impairs growth, despite the presence of alternative, less efficient uptake systems (Fuentes et al., 2013). To evaluate the impact of these deletions, we compared the growth of parental MDO and MGX strains in M9 minimal medium supplemented with either glucose or mannose as sole carbon source (Table 2).

**Table 2:** Growth rates (μ) and specific uptake rates (qs) for the studied *E. coli* strains grown on glucose and mannose.

| Strains | Glucose |  | Mannose |  |
| --- | --- | --- | --- | --- |
| | $\mu$ ($\text{h}^{-1}$) | $q_s$<br>($\text{mmol.g}_{\text{cdw}}^{-1}.\text{h}^{-1}$) | $\mu$ ($\text{h}^{-1}$) | $q_s$<br>($\text{mmol.g}_{\text{cdw}}^{-1}.\text{h}^{-1}$) |
| MDO | $0.47 \pm 0.04$ | $6.14 \pm 1.02$ | $0.26 \pm 0.01$ | $2.05 \pm 0.09$ |
| MGX | $0.15 \pm 0.01$ | $1.11 \pm 0.20$ | $0.10 \pm 0.01$ | $1.31 \pm 0.07$ |
| MGX1 | $0.26 \pm 0.02$ | $5.27 \pm 1.34$ | $0.20 \pm 0.01$ | $1.90 \pm 0.09$ |
| PFKA1 | $0.20 \pm 0.01$ | $2.73 \pm 0.43$ | $0.14 \pm 0.01$ | $1.09 \pm 0.06$ |

As expected, the MGX strain grew roughly three times more slowly on both sugars. To restore sugar uptake and growth, we introduced a plasmid (pWKS) expressing the *galP* gene, encoding *E. coli* galactose permease under the control of the Plac promoter, into the MGX strain, generating MGX1. Consistent with previous reports, induction of recombinant GalP partially restored glucose uptake and growth (Hernández-Montalvo et al., 2003): on glucose (mannose), the growth rate of the MGX1 strain was roughly 55% (75%) of the MDO strain’s (Table 2).

### 3.3 Deletion of phosphofructokinase A leads to accumulation of intracellular hexose-phosphate

#### Deletion of phosphofructokinase A leads to accumulation of intracellular hexose-phosphate

Inactivation of the PTS leads to the accumulation of unphosphorylated monosaccharides in the cytoplasm, making them available as substrates for GPs involved in reverse-phosphorolysis. To further elevate intracellular levels of hexose-1-phosphates, the glycosyl donor substrates required by GPs, we deleted the *pfkA* gene, which encodes the glycolytic enzyme phosphofructokinase A. This enzyme catalyzes the conversion of fructose-6-phosphate (F6P) to fructose-1,6-bisphosphate (FBP). Deletion of *pfkA* is known to result in the intracellular accumulation of glucose-6-phosphate (G6P) and F6P [42], which should in turn increase the availability of glucose-1-phosphate (G1P) and mannose-1-phosphate (M1P) for GP-mediated synthesis.

### Restoration of growth in the PFKA strain via GalP expression

The PFKA strain, generated by deleting *pfkA* from the MGX1 strain, showed no detectable growth after three days in minimal medium with glucose or mannose as sole carbon source. Introduction of a plasmid (pWKS) expressing the galactose permease gene (*galP*), yielding the PFKA1 strain, restored glucose uptake and partially recovered growth in minimal medium with glucose (growth rate: 0.20 ± 0.01 h⁻¹, less than half that of the MDO strain; see Table 2). On mannose, PFKA1 had a growth rate of 0.14 ± 0.01 h⁻¹, indicating viability, albeit with slower growth.

### Intracellular metabolite profiles of the MDO, MGX1, and PFKA1 strains on glucose and mannose

The results of metabolomic analyses of MDO, MGX1, and PFKA1 strains grown on either glucose or mannose as sole carbon source are presented in Figure 2.

**Figure 2:**
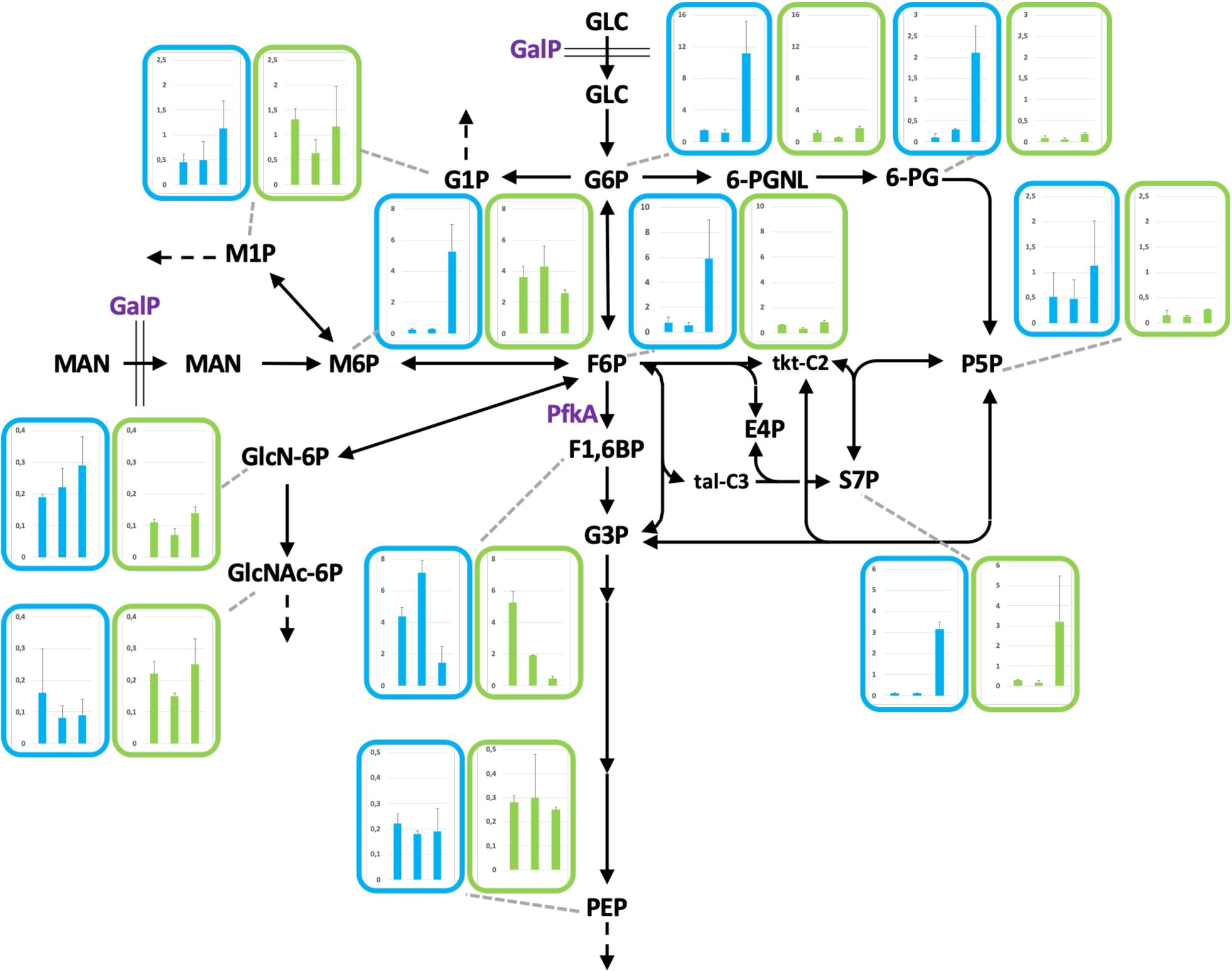
Absolute intracellular concentrations of phosphorylated sugars in strains grown with glucose or mannose as the sole carbon sources. Values are expressed in µmol·gDW^−1^ and represent the mean of two independent experiments; standard deviations are indicated as error bars. Data from glucose- and mannose-based cultures are shown in blue and green boxes, respectively. In each box, the left bar corresponds to the MDO strain, the middle bar to strain MGX1, and the right bar to strain PFKA1. *Abbreviations: GalP, galactose permease; PfkA, phosphofructokinase I; Metabolites: GLC, glucose; Man, mannose; G1P, glucose-1-phosphate; M6P, mannose-6-phosphate; F6P, fructose-6-phosphate; G6P, glucose-6-phosphate; M1P, mannose-1-phosphate; F1,6BP, fructose-1,6-bisphosphate; 6-PGNL, 6-phosphogluconolactone, 6-PG, 6-phosphogluconate; P5P, pentose-5-phosphate; E4P, erythrose-4-phosphate; S7P, sedoheptulose-7-phosphate; tkt-C2, C2 fragment bound to transketolase ; tal-C3, C3 fragment bound to transaldolase; G3P, glyceraldehyde-3-phosphate; PEP, phosphoenolpyruvate; GlcN-6P, glucosamine-6-phosphate; GlcNac-6P, N-acetylglucosamine-6-phosphate*.

On glucose (Figure 2, blue box), the intracellular metabolite profiles of MGX1 were similar to those of MDO, as expected, since deletion of the PTS transporter in MGX1 is not expected to significantly impact central carbon metabolism. In contrast, the profiles of PFKA1 deviated significantly from those of the two other strains. Most notably, the intracellular concentration of fructose-1,6-bisphosphate (FBP) was twofold lower than in MDO and fivefold lower than in MGX1. This is consistent with deletion of the *pfkA* gene, since FBP is the direct product of the phosphofructokinase-catalyzed reaction. Correspondingly, concentrations of the upstream glycolytic intermediates glucose-6-phosphate (G6P) and fructose-6-phosphate (F6P) levels were approximately 10- and 8-fold higher than in MDO, respectively. Increases were also observed for other hexose-phosphates, including a 20-fold increase in the concentration of mannose-6-phosphate (M6P), and a strongly elevated combined pool of glucose-1-phosphate (G1P) and mannose-1-phosphate (M1P), which are not singularly distinguishable by ion chromatography. In addition, the PFKA1 strain had elevated levels of several pentose phosphate pathway (PPP) intermediates, such as 6-phosphogluconate (6-PG), pentose-5-phosphates (P5P; including ribose-5-phosphate and ribulose-5-phosphate), and in particular, sedoheptulose-7-phosphate (S7P), whose concentration was 10–20-fold higher than in MDO and MGX1. These results point to a broad re-routing of carbon fluxes. Indeed, previous studies have shown that *pfkA* deletion induces a metabolic shift from glycolysis toward the PPP (Hollinshead et al., 2016), which explains the observed accumulation of PPP-related metabolites.

Similar trends in metabolite concentrations were observed when the same strains were grown on mannose as sole carbon source (Figure 2, green box). However, a major exception was the pronounced accumulation of M6P in both the MDO and MGX1 strains, as expected, given that M6P is the first phosphorylated intermediate in mannose metabolism. In the PFKA1 strain, FBP levels were approximately tenfold lower than in MDO, consistent with the altered metabolic flux associated with the absence of PfkA. No general increase in PPP intermediates was observed however, with the exception of S7P, whose elevation remains unexplained at this stage. Notably, the accumulation of hexose-phosphates in the PFKA1 strain was much less pronounced during growth on mannose than during growth on glucose. While M6P and M1P levels remained similar, G6P and F6P were slightly higher than in the MDO strain (F6P, 0.88 ± 0.11 vs. 0.64 ± 0.05; G6P, 1.75 ± 0.21 vs. 1.13 ± 0.34). Surprisingly, intracellular concentrations of several metabolites, including the G1P/M1P pool, were lower in MGX1 than in MDO, despite manB overexpression. Under mannose conditions therefore, PFKA1 accumulated higher levels of these intermediates than MGX1.

In summary, these results show that PFKA1 has an intracellular metabolite profile that differs from those of MDO and MGX1, with a strong accumulation of hexose-phosphates and reduced FBP levels on glucose. These differences are less pronounced on mannose, where only FBP remains consistently low. Given MGX1’s unexpectedly lower metabolite levels on mannose, the increased accumulation of GP substrates under mannose conditions evidenced here in PFKA1 make it the most promising candidate for mannooligosaccharide production.

### 3.6 Strain PFKA1 produces β-1,2-mannobiose *in vivo* by GP-mediated synthesis

β-1,2-mannobiose production was evaluated in the double mutant PFKA1 strain lacking both PTS and PfkA. This disaccharide can be synthesized by the glycoside-phosphorylase Teth514_1788 (Ahmadipour et al., 2023), a member of the GH130 family of carbohydrate-active enzymes (CAZy database, http://www.cazy.org/ (Drula et al., 2022)). The PFKA1 strain was transformed with plasmid pBAD harboring Teth514_1788 under arabinose-inducible control. The resulting strain, PFKA1-Teth514_1788, was cultivated along with the parental PFKA1 strain as a control in baffled shake flasks containing minimal medium with mannose as the sole carbon source. NMR spectra of the culture supernatants are shown in Figure 3.

**Figure 3:**
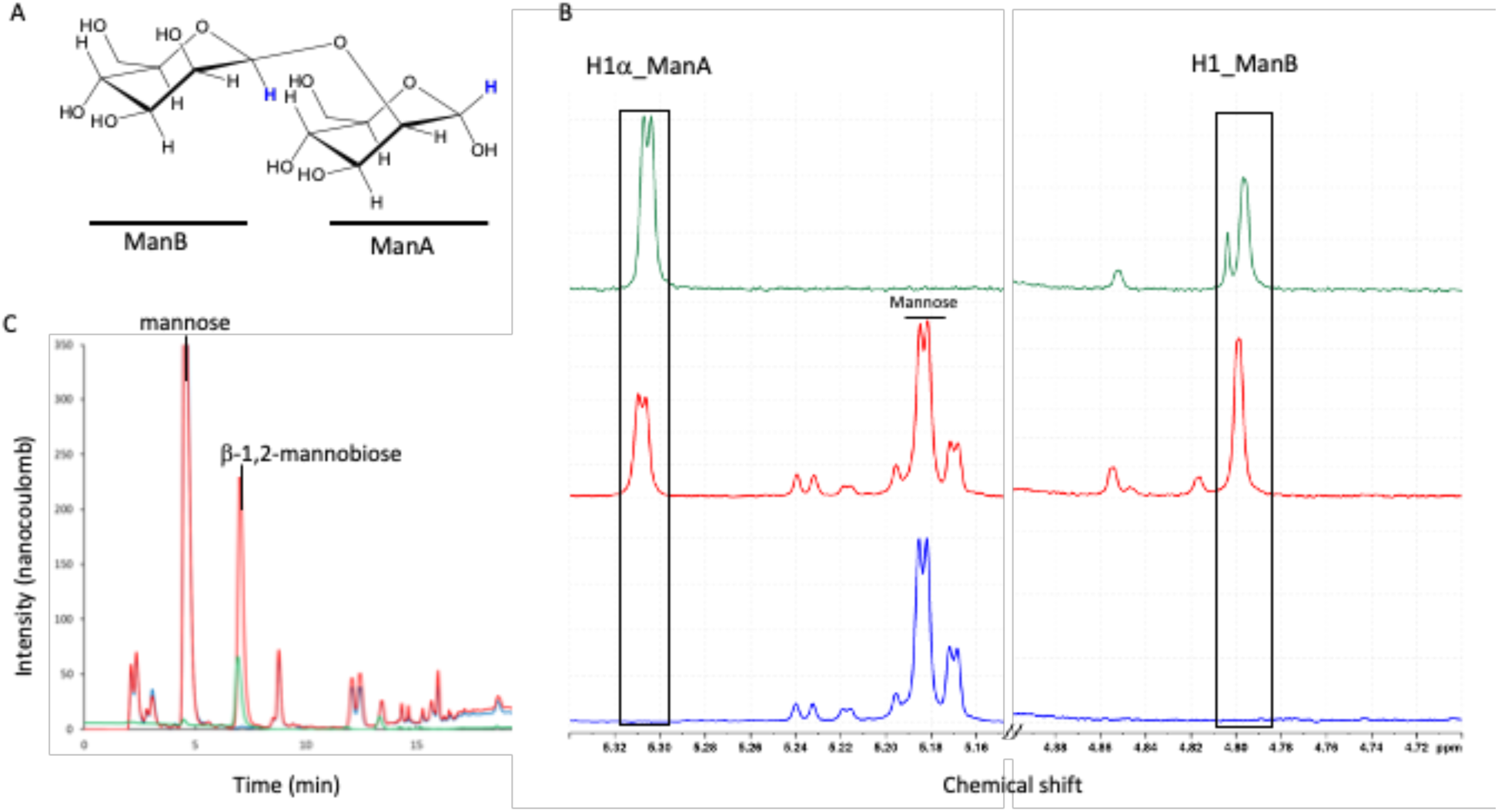
A. Chemical structure of β-1,2-mannobiose, showing the anomeric protons of the reducing (ManA) and the non-reducing (ManB) mannosyl moieties (highlighted in blue), corresponding to the signals observed in the NMR spectra in panel B. B. Selected regions of the 1D ^1^H NMR spectra of β-1,2-mannobiose corresponding to the anomeric proton signals. C. HPAEC-PAD chromatograms showing that the β-1,2-mannobiose produced in the culture supernatant co-eluted with a β-1,2-mannobiose standard. *Bright green lines, β-1,2-mannobiose standard; red, culture supernatant of PFKA1-Teth514_1788; blue, culture supernatant of PFKA1-control*.

Two well-defined peaks were detected in the spectra of the culture supernatants of the PFKA1-Teth514_1788 strain that were absent from the control samples (Figure 3B), and whose intensity increased over the course of cultivation (data not shown), suggesting progressive secretion of the product into the medium. These signals correspond to those observed in the NMR spectrum of β-1,2-mannobiose, as previously annotated (Shibata et al., 1993). Based on their chemical shifts, the two peaks at 5.31 ppm and 4.79 ppm correspond to the α-anomeric proton (H1α) of the reducing mannosyl moiety (ManA) and to the anomeric proton of the non-reducing mannosyl moiety (ManB) respectively (Figure 3A). These chemical shift assignments were further corroborated by high-performance anion-exchange chromatography with pulsed amperometric detection (HPAEC-PAD), which confirmed the presence of β-1,2-mannobiose in the culture supernatant through elution at the same retention times as the standard (Figure 3C).

### 3.7 GP-based production of β-1,4-mannobiose and laminaribiose

The versatility of our engineered strain was tested through the production of two additional oligosaccharides, β-1,4-mannobiose and laminaribiose. For β-1,4-mannobiose, the PFKA1 strain was transformed with the plasmid pBAD-UhgbMP, encoding a β-1,4-mannoside-phosphorylase (Ladevèze et al., 2013). Presence of β-1,4-mannobiose in the culture supernatant was confirmed by NMR and HPAEC-PAD, with characteristic signals matching those of a commercial standard (Figure S1), demonstrating efficient production and extracellular accumulation of the disaccharide.

Laminaribiose synthesis was catalyzed by the laminaribiose-phosphorylase ACL0729 using glucose and G1P as substrates (Nihira et al., 2012). Figure S2B presents an expanded view of the spectral regions spanning 5.28–5.20 ppm and 4.76–4.67 ppm, where distinct signals are observed exclusively in the supernatant of the *E. coli* PFKA1-pBAD-ACL0729 strain and in a commercial laminaribiose standard, but not in the supernatant of the untransformed PFKA control strain. Spiking the PFKA1-pBAD-ACL0729 supernatant with commercial laminaribiose resulted in a clear increase in the same signals, confirming the presence of the compound. These NMR results were validated by high-performance anion-exchange chromatography with pulsed amperometric detection (HPAEC-PAD) (Figure S2C).

These results confirm that the proposed PFKA chassis strain is compatible with multiple GPs from different CAZy families and can efficiently synthesize and secrete diverse β-linked oligosaccharides *in vivo*, highlighting its potential as a platform for tailored oligosaccharide biosynthesis.

### 3.8 Synergistic effect of PTS and *pfkA* deletions on β-1,2-mannobiose production

After demonstrating mannobiose production in strain PFKA1 as proof of concept, we next aimed to assess the individual and combined contributions of the underlying genetic modifications by comparing β-1,2-mannobiose production in different strains. The pBAD-Teth514_1788 plasmid was introduced into strains MDO, MGX1, and PFKA1, which were then cultivated in bioreactors under previously described conditions using 10 g·L^−1^ mannose as the sole carbon source. NMR analyses of the supernatants at the end of cultivation are summarized in Table 3.

**Table 3:**
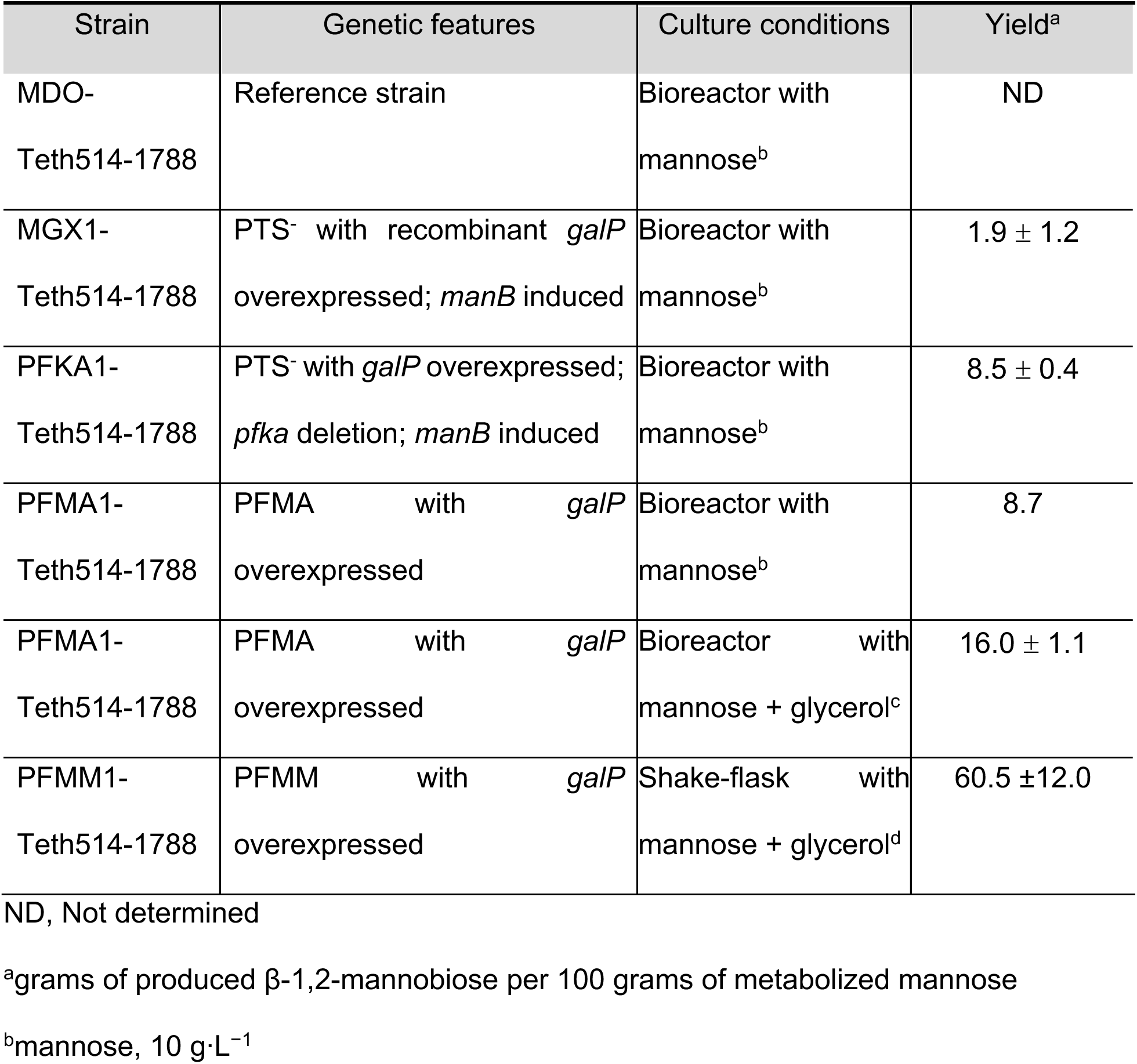
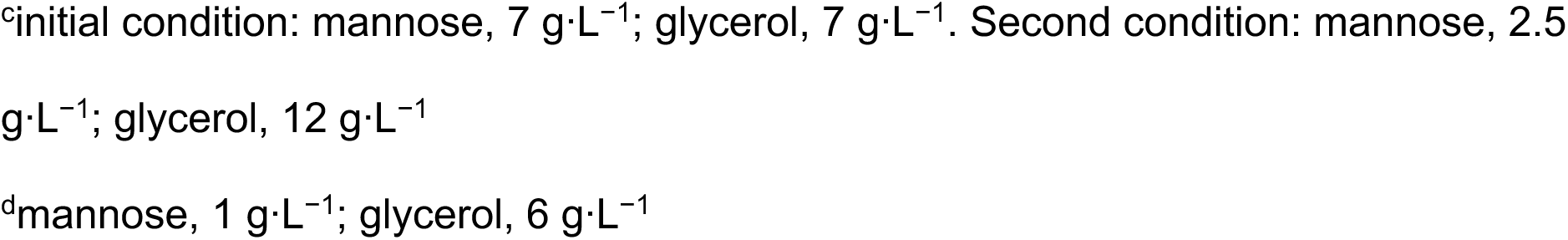
β-1,2-mannobiose production yields of engineered strains under different culture conditions.

| Strain | Genetic features | Culture conditions | Yield<sup>a</sup> |
| --- | --- | --- | --- |
| MDO-Teth514-1788 | Reference strain | Bioreactor with mannose<sup>b</sup> | ND |
| MGX1-Teth514-1788 | PTS<sup>-</sup> with recombinant *galP* overexpressed; *manB* induced | Bioreactor with mannose<sup>b</sup> | 1.9 ± 1.2 |
| PFKA1-Teth514-1788 | PTS<sup>-</sup> with *galP* overexpressed; *pfka* deletion; *manB* induced | Bioreactor with mannose<sup>b</sup> | 8.5 ± 0.4 |
| PFMA1-Teth514-1788 | PFMA with *galP* overexpressed | Bioreactor with mannose<sup>b</sup> | 8.7 |
| PFMA1-Teth514-1788 | PFMA with *galP* overexpressed | Bioreactor with mannose + glycerol<sup>c</sup> | 16.0 ± 1.1 |
| PFMM1-Teth514-1788 | PFMM with *galP* overexpressed | Shake-flask with mannose + glycerol<sup>d</sup> | 60.5 ± 12.0 |

ND, Not determined
<sup>a</sup>grams of produced β-1,2-mannobiose per 100 grams of metabolized mannose
<sup>b</sup>mannose, 10 g·L<sup>-1</sup>
<sup>c</sup>initial condition: mannose, 7 g·L<sup>-1</sup>; glycerol, 7 g·L<sup>-1</sup>. Second condition: mannose, 2.5 g·L<sup>-1</sup>; glycerol, 12 g·L<sup>-1</sup>
<sup>d</sup>mannose, 1 g·L<sup>-1</sup>; glycerol, 6 g·L<sup>-1</sup>

No β-1,2-mannobiose was detected in the supernatant of the MDO strain, indicating that in the presence of an intact PTS, mannose is internalized exclusively in its phosphorylated form, precluding the reverse-phosphorylation reaction required for mannobiose synthesis. Strain MGX1, lacking the PTS, produced a detectable amount of mannobiose (0.4 ± 0.2 mM) (Figure S3). This result highlights the pivotal role of PTS deletion in enabling the import of unphosphorylated mannose into the cytoplasm—a prerequisite for the synthetic mannobiose production. Furthermore, overexpression of the *manB* gene in MGX1 may result in increased intracellular availability of M1P, resulting from *manB*-driven conversion of M6P to M1P.

The PFKA1 strain, featuring both PTS and *pfkA* deletions along with *manB* overexpression, had the highest β-1,2-mannobiose titer (1.8 mM ± 0.1, corresponding to 0.62 g·L⁻¹) and a conversion yield of 8.5 ± 0.4 %, an almost four-fold improvement over strain MGX1 (Figure S3). This increase is likely due to elevated intracellular concentrations of M1P resulting from the redirected carbon flux caused by *pfkA* deletion. In summary, the fact that strains with an intact PTS system fail to produce detectable amounts of mannobiose show that PTS deletion is crucial for β-1,2-mannobiose biosynthesis. Two additional engineering effects, enhanced mannose transport and increased intracellular M1P availability, further boost β-1,2-mannobiose yields in the PFKA1 strain.

### 3.9 Further chassis improvement

#### 3.9.1 Elimination of an acetylated sugar contaminant *via maa* gene deletion

The NMR spectra of supernatants from strain PFKA1 cultivated in minimal medium contained an un unexpected peak, which could not be assigned to any known sugar standards. Preliminary spectral analysis suggested that the unknown compound was an acetylated monosaccharide. Given the broad substrate specificity of maltose O-acetyltransferase (encoded by *maa*), it was hypothesized that this enzyme catalyzes the acetylation of internalized monosaccharides with acetyl-CoA as an acetyl donor—a reaction that is presumably facilitated by the unique metabolic configuration of the PFKA1 strain, whose defective PTS system allows the uptake of unphosphorylated sugars into the cytosol.

The *maa* gene was therefore knocked out, both to confirm the origin of this acetylated contaminant and to prevent a potentially competitive side reaction diverting precursors from β-1,2-mannobiose synthesis. Two strains, PFKA1 and its isogenic Δ*maa* derivative (PFMA1), both harboring the plasmid pBAD-Teth514_1788, were cultivated in M9 minimal medium in shake-flasks supplemented with 6 g·L⁻¹ mannose. The extracellular metabolite profiles were analyzed by NMR (Figure S4).

Deletion of *maa* led to the complete disappearance from the culture supernatant of the acetylated sugar contaminant, which we tentatively identified as 6-O-acetyl-mannose (6-O-AcMan). However, this deletion did not result in a measurable increase in β-1,2-mannobiose yield compared to the PFKA1 strain (see Table 3).

#### 3.9.2. Improved mannose conversion through glycerol co-feeding and partial relief of carbon catabolite repression

Maximizing the conversion efficiency of mannose is particularly important given its relatively high cost as a carbon source. In the proof-of-concept configuration, mannose serves both as a carbon source for cellular growth and as a substrate for β-1,2-mannobiose biosynthesis, which intrinsically constrains the maximum achievable yield. However, the ability of PTS-deficient *E. coli* strains to co-utilize substrates offers a potential means to decouple biomass formation from product synthesis. As well as mediating sugar uptake, the PTS plays a key regulatory role in carbon catabolite repression (CCR), and its disruption enables simultaneous assimilation of multiple carbon sources. Glycerol was selected as a co-substrate due to its efficient uptake kinetics in the PTS-deficient background and its low cost (relative to mannose). To evaluate this strategy, a first bioreactor cultivation was carried out using strain PFMA1 harboring the pBAD-Teth514_1788 plasmid in minimal medium supplemented with equimolar concentrations of mannose and glycerol (7 g·L⁻¹ each) (Figure S5A).

As anticipated, glycerol was rapidly consumed within the first 25 h of cultivation, whereas mannose uptake was incomplete, with approximately 75% of the initial concentration remaining unutilized. Importantly though, and as expected, mannose was co-consumed, presumably because of the alleviation of CCR in this strain. The final concentration of β-1,2-mannobiose was 0.70 mM; however, the mannose-to-β-1,2-mannobiose conversion yield was significantly improved, reaching 16.8% compared with 9% in the mannose-only setup. These results were corroborated in a second bioreactor experiment in which the glycerol concentration was doubled and the mannose concentration reduced to one-third of its initial level (Figure S5B). Under these conditions, mannose utilization was also incomplete, with approximately 50% of the initial concentration remaining unutilized. However, the mannose-to-β-1,2-mannobiose conversion yield was similar to that observed previously, reaching 15.2%. Overall, under mannose–glycerol co-feeding conditions, the mannose-to-β-1,2-mannobiose conversion yield reached 16.0 ± 1.1% (see table 3).

#### 3.9.3. Decoupling growth and production to increase mannose conversion into β-1,2-mannobiose

These preliminary results support the feasibility of decoupling growth and production by co-substrate utilization. Nonetheless, there remains substantial room for optimization, as these data indicate that approximatively 20% of the internalized mannose was directed toward β-1,2-mannobiose synthesis, with the remainder (> 80%) still funneled into central carbon metabolism.

To further decouple cellular growth from β-1,2-mannobiose biosynthesis we deleted the *manA* gene, generating strain PFMM1. As illustrated in Figure 4, deletion of *manA* creates a “closed circuit” by preventing the conversion of M6P into F6P, thereby channeling M6P preferentially towards β-1,2-mannobiose synthesis. The potential flux diversion from M1P to GDP-mannose *via* M1P guanylyl-transferase (encoded by *CpsB*) is likely negligible. In the PFMM1 strain, deletion of *wcaJ* abolishes colanic acid biosynthesis, for which GDP-mannose is an essential precursor. Cultivations were carried out in M9 minimal medium under standard conditions in shake-flasks with glycerol (6 g·L^−1^) and mannose (1 g·L^−1^) as carbon sources.

**Figure 4.**
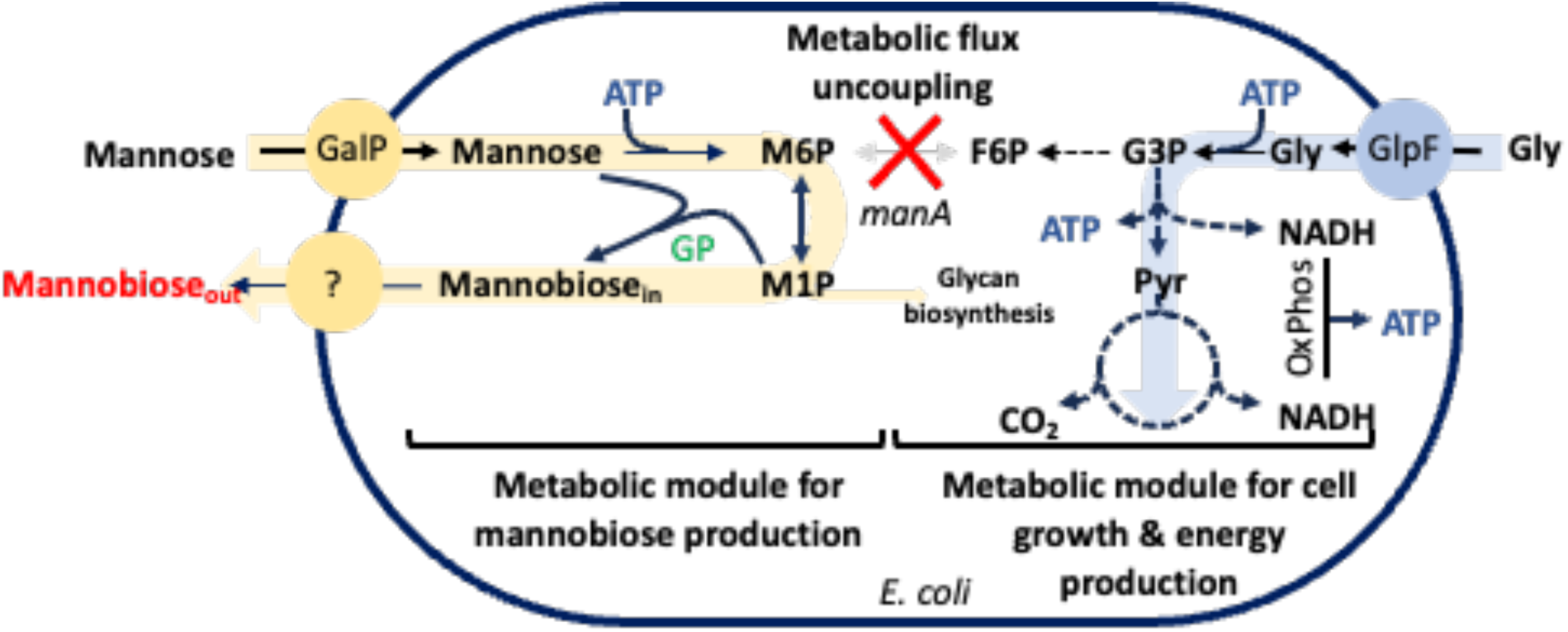
β-1,2-mannobiose production *via* co-substrate metabolism enabled by metabolic uncoupling of growth and product synthesis through the deletion of *manA*. *GalP, galactose permease; manA, phosphomannose isomerase; GP, glycoside-phosphorylase; M6P, mannose-6-phosphate; F6P, fructose-6-phosphate; M1P, mannose-1-phosphate; GDP-Man, GDP-mannose; G3P, glycerol-3-phosphate; Pyr, pyruvate; ATP, adenosine-triphosphate; Gly, glycerol; OxPhos, oxidative phosphorylation*.

The β-1,2-mannobiose concentration at the end of cultivation was 0.84 ± 0.07 mM, corresponding to a mannose conversion rate of 60.5 ± 12%, substantially higher than in the *manA-*positive strain (16%) (Table 3). Cultivating this PFMM1 strain in a bioreactor led to a final β-1,2-mannobiose titer of 1.21 mM and a conversion rate of 58 %, consistent with flask-scale experiments, and demonstrating the robustness and scalability of the process (data not shown).

## 4. Conclusion

In this study, we engineered an *E. coli* strain for the biosynthesis of disaccharides *via* GPs, leveraging a dual metabolic blockade strategy: inactivation of the PTS, allowing import of non-phosphorylated monosaccharides, and deletion of the *pfkA* gene, leading to intracellular accumulation of the phosphorylated sugar precursors (M1P, G1P) required for synthesis.

Our results show that mannose uptake in PTS⁻ strains is restored by heterologous expression of the *galP* permease, enabling partial growth on mannose. Introduction of the Δ*pfkA* mutation resulted in marked accumulation of intracellular phosphohexoses, particularly under glucose conditions, but also to a lesser extent under mannose, establishing strain PFKA1 as a favorable metabolic chassis for oligosaccharide biosynthesis.

Through expression of glycoside-phosphorylases from distinct CAZy families (GH130 and GH 94), we achieved *in vivo* production of two mannobioses (β-1,2 and β-1,4-mannobiose) and laminaribiose, thus demonstrating the modularity and versatility of the PFKA chassis for tailored oligosaccharide synthesis. The combination of PTS deletion, *manB* overexpression, and *pfkA* knockout had a pronounced synergistic effect, leading to a mannose-to-β-1,2-mannobiose conversion efficiency of approximatively 9% in the PFKA1 strain, far higher than measured in the corresponding *pfkA*-positive strain (2%).

We also observed the accumulation of an acetylated form of mannose in the culture supernatant, which disappeared upon deletion of the *maa* gene. This gene encodes maltose O-acetyltransferase, an enzyme known for its broad sugar substrate specificity (Brand and Boos, 1991). To our knowledge, the accumulation of acetylated mannose in culture supernatants has never previously been reported, and likely results from the import of unphosphorylated mannose under PTS-deficient conditions.

Finally, we demonstrated that decoupling biomass formation from product synthesis through co-substrate feeding is an effective strategy to improve mannose-to-β-1,2-mannobiose conversion in engineered *E. coli*. The PTS-deficient background enables simultaneous assimilation of glycerol and mannose, with glycerol supporting cellular energy demands and mannose being redirected toward disaccharide biosynthesis. While co-feeding only modestly improved titers, the conversion yield nearly doubled compared with mannose-only cultures, highlighting the benefit of alleviating CCR. Further enhancement was achieved by deleting *manA*, thereby blocking the competing metabolic branch from M6P to central carbon metabolism. This modification significantly increased mannose utilization in β-1,2-mannobiose synthesis, with conversion rates of nearly 60% in both shake-flask and bioreactor cultures. Collectively, these results validate the effectiveness of growth-production decoupling strategies and metabolic channeling to optimize the biosynthesis of oligosaccharides in microbial systems.

Overall, our approach illustrates the power of rational *E. coli* metabolic engineering combined with GP-based catalysis for the sustainable biosynthesis of costly sugars, with potential applications in prebiotics and in the cosmetic and pharmaceutical industries.

## 5. Conflicts of interest

PT, JD, GPV, and FL declare competing financial interest related to the primary production process which is covered by a patent application filed under the PCT(WO/2021/229185) and licensed to Sweetech. JD is a co-founder of Sweetech SAS. LT declares no competing interests.

## 6. Funding

This work was supported by the Agence Nationale de la Recherche [grant number ANR-16-CE20-0006-03] and by Toulouse Tech Transfer, a technology transfer acceleration company, with the mission to transfer and commercialize innovative technologies developed by the research laboratories of the west Occitanie region (http://www.toulouse-tech-transfer.com).

## 7. Acknowledgement

The authors thank MetaToul (Metabolomics & Fluxomics Facilities, Toulouse, France, www.metatoul.fr) and its staff for technical support and access to the MS and NMR facilities. MetaToul is part of the French National Infrastructure for Metabolomics and Fluxomics (www.metabohub.fr), funded by the Agence Nationale de la Recherche under the France 2030 program (MetaboHUB ANR-11-INBS-0010 ; MetEx+ ANR-21-ESRE-0035; MetaboHUB (JVCE) ANR-24-INBS-0012). The authors also thank the PICT-ICEO facility dedicated to enzyme screening and discovery, part of the Integrated Screening Platform of Toulouse (PICT, IBiSA) for providing access to HPAEC-PAD and enzyme purification equipment. PICT-ICEO is a member of IBISBA-FR (https://doi.org/10.15454/08BX-VJ91), the French node of the European research infrastructure, IBISBA (www.ibisba.eu). The authors also thank Dr Éric Samain for his valuable support and insightful contributions.

## 8. Author contributions

PT: Conceptualization, Methodology, Formal analysis, Investigation, Writing - Original Draft. JD: Conceptualization, Methodology, Formal analysis, Investigation. LT: Methodology. GPV: Conceptualization, Validation, Writing - Review & Editing, Supervision, Funding acquisition. FL: Conceptualization, Validation, Writing- Review and Editing, Visualization, Supervision, Project administration, Funding acquisition. All authors have read and approved the final version of the manuscript.

