## Supplemental Table and Figures for "Metabolic engineering of *Escherichia coli* strains for the *in vivo* synthesis of GP-mediated oligosaccharides"

-

Pietro Tedesco#, Julien Durand#, Laurence Tarquis, Gabrielle Potocki-Veronese*, Fabien Létisse*

### Equally contributed to the work

* Co-corresponding authors

Fabien Létisse :

Gabrielle Potocki-Veronese:

FL: Institut de Pharmacologie et de Biologie Structurale (IPBS), Université de Toulouse, CNRS, UPS, Toulouse, France

JD, LT, PT et GPV: TBI, Université de Toulouse, CNRS, INRAE, INSA, 31077 Toulouse, France

PT (Current address): Stazione Zoologica Anton Dohrn, Department of Ecosustainable Marine Biotechnology, Naples, Italy

JD (Current address): Sweetech SAS, INSA, TBI, 135 Av. de Rangueil, 31077 Toulouse, France

**Table S1. Oligonucleotide sequences of the primers used for gene deletions**

| *Target gene* | *Name* | *Sequence 5’ – 3’* |
|---|---|---|
| *ptsG* upstream | P1 | *f* CTAGGATCCGCCGAAAATTGGGCGGTGAATAAC |
| | P2 | *r* GTAAAGCTTCTGGTAAGCAGCCCACTGAGAGAAG |
| *ptsG* downstream | P3 | *f* CTCAAGCTTGCGCCGATCCTGTACATCATCC |
| | P4 | *r* CGTGTCGACCGTTAATCCTAATCTGCCACGCACC. |
| *manXYZ* upstream | P5 | *f* ACTCGAGAATGGCGATGAAGACCAG |
| | P6 | *r* CCAGTGCCACCAGTTCATC |
| *manXYZ* downstream | P7 | *f* CAAGCTTCGATTCCGGAAGTGGTGAC |
| | P8 | *r* AGGATCCATGCCCCCATGACAAACAG |
| *pfkA* upstream | P9 | *f* GGATCCAGGCGTCGGGGTTATCGTG |
| | P10 | *r* AAGCTTCGGTCTTCATACAGACCCAGATAGC |
| *pfkA* downstream | P11 | *f* AAGCTTGGCCATTGCCGGTGGCTGTG |
| | P12 | *r* CTCGAGCCACCGTGTGACTGACGAATC |
| *maa* | P13 | *f* ATGAGCACAGAAAAAGAAAAGATGATTGCTGGTGAGTTGTATCGC TCGGCGTGTAGGCTGGAGCTGCTTC |
| | P14 | *r* TTACAATTTTTTAATTATTCTGGCTGGATTACCGCCCACGACAACG TTGTCATATGAATATCCTCCTTAG |
| *manA* | P15 | *f* ATTATGCGCAGCACAGCCACTCTCCATTCAGGTTCATCCAAACAA ACACAGTGTAGGCTGGAGCTGCTTC |
| | P16 | *r* ACACGCGCTAAACGGCCGTGGCCTTTGACAGTCACCGGTGATTC GTTGGCCATATGAATATCCTCCTTAG |
| *GalP* cloning | P17 | *f* TTGTCGACTTAAGGAGGGCATCATGCCTGAC |
| | P18 | *r* GTTCTAGATGACTGCAAGAGGTGGCTTCC |
| *Uhgb_MP* cloning | P19 | *f* CACCATGAGTATGAGTAGCAAAGTTATT–3′ |
| | P20 | *r* GATGATGCTTGTACGTTTGGTAAATTC |

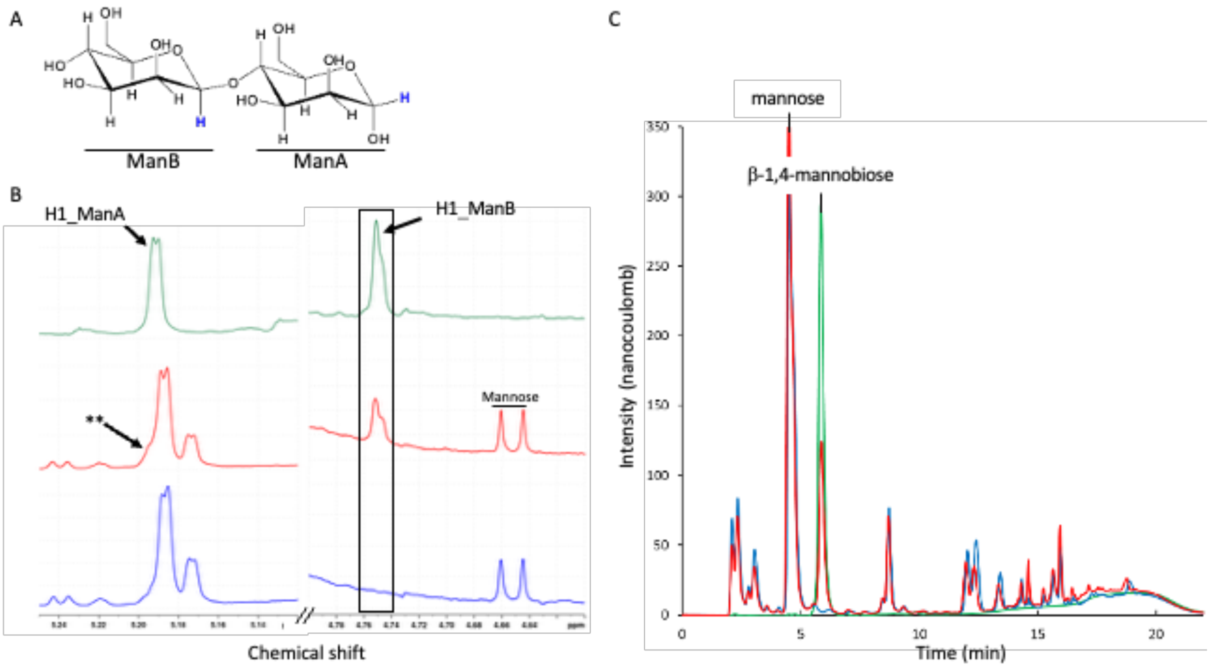


Figure S1: A. Chemical structure of β-1,4-mannobiose, showing the anomeric protons of the reducing (ManA) and the non-reducing (ManB) mannosyl moieties (highlighted in blue), corresponding to the signals observed in the NMR spectra in panel B. B. Selected regions of the 1D $^{1}$H NMR spectrum of β-1,4-mannobiose corresponding to the anomeric proton signals (**, slight shoulder). C. HPAEC-PAD chromatograms showing that the β-1,4-mannobiose produced in the culture supernatant and co-elute with the commercial β-1,4-mannobiose standard.

*Green, commercial standard; red, culture supernatant of PFKA1-pBAD-UhgbMP; blue, culture supernatant of PFKA1-control.*

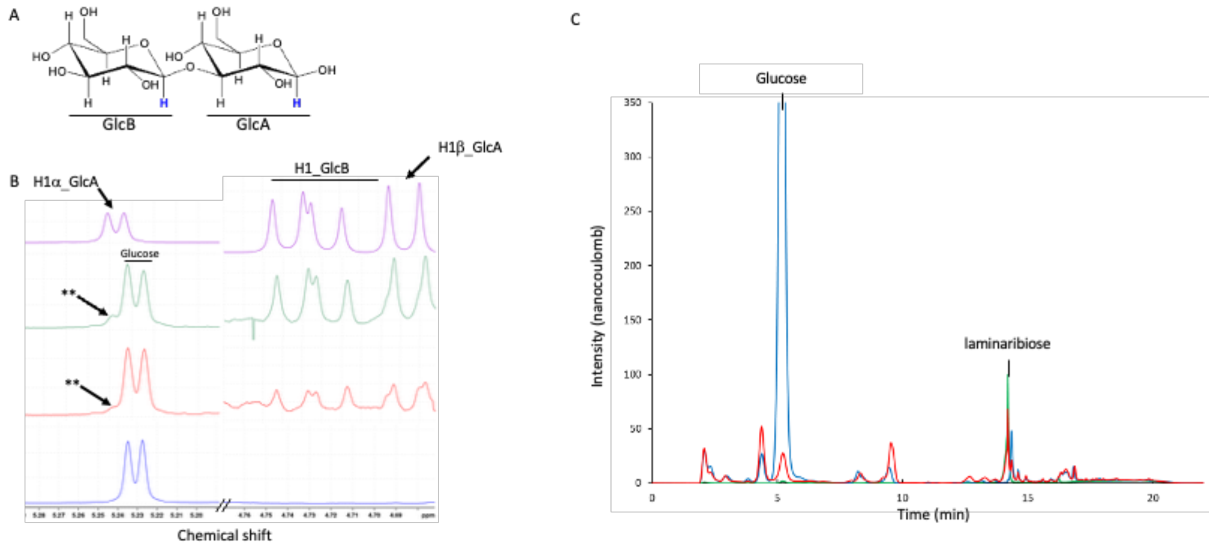


Figure S2: A. Chemical structure of laminaribiose (β-D-gluco-hexopyranosyl-(1->3)-β-D-gluco-hexopyranose), showing the anomeric protons of the reducing (GlcA) and the non-reducing (GlcB) glucosyl moieties (highlighted in blue), corresponding to the signals observed in the NMR spectra in panel B. B. Selected regions of the 1D $^{1}$H NMR spectrum of laminaribiose corresponding to the anomeric proton signals (**, slight shoulder). C. HPAEC-PAD chromatograms showing that the laminaribiose produced in the culture supernatant and co-elute with the commercial laminaribiose.

*Green, commercial standard; brown, culture supernatant of PFKA1-pBAD-ACL0729 spiked with commercial laminaribiose; red, culture supernatant of PFKA1-pBAD-ACL0729 ; blue, culture supernatant of PFKA1-control.*

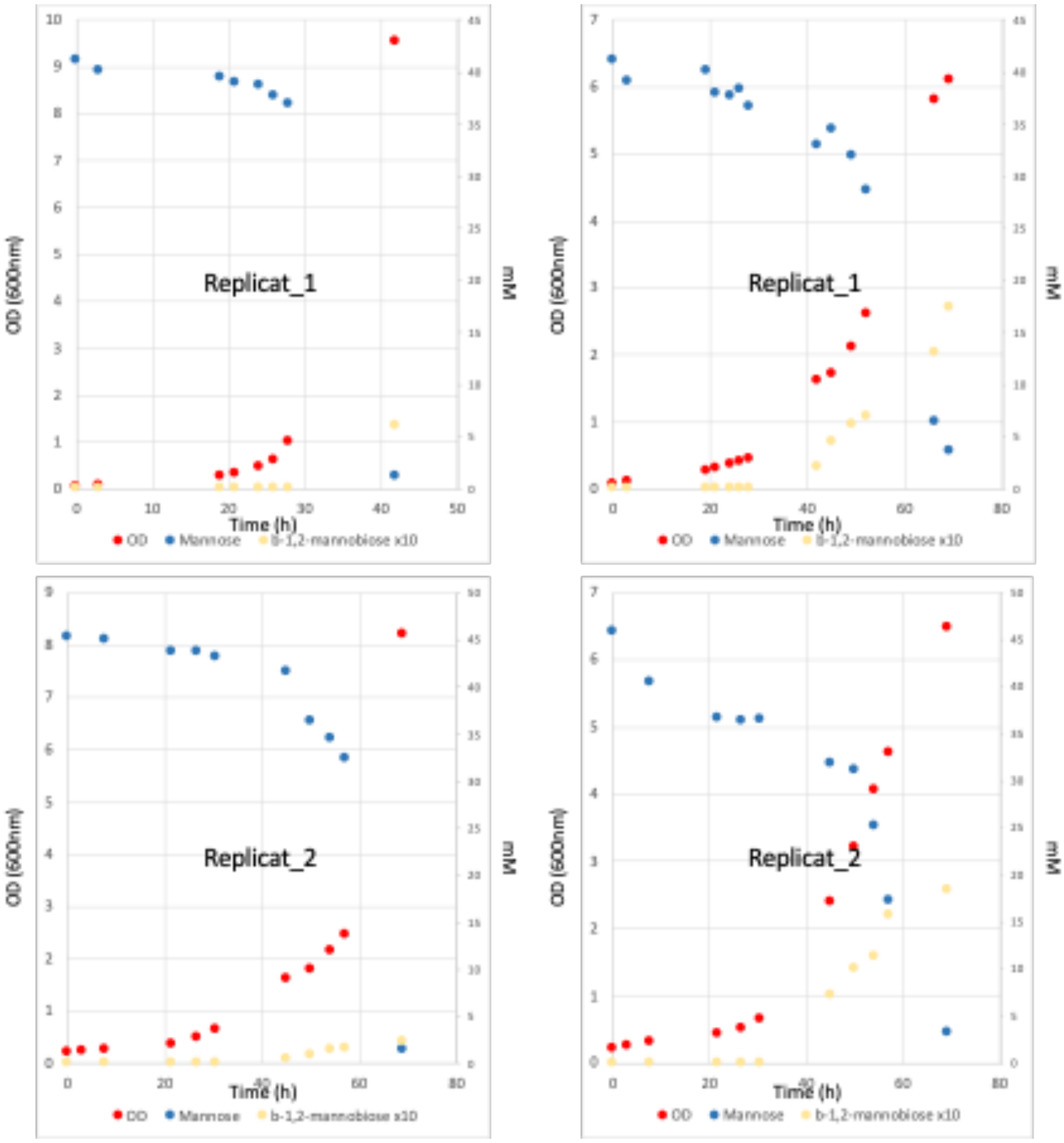


Figure S3. Growth profiles of E. coli MGX-Teth514-1788 (left) and PFKA1-Teth514_1788 (right) strains cultivated in bioreactors on mannose.

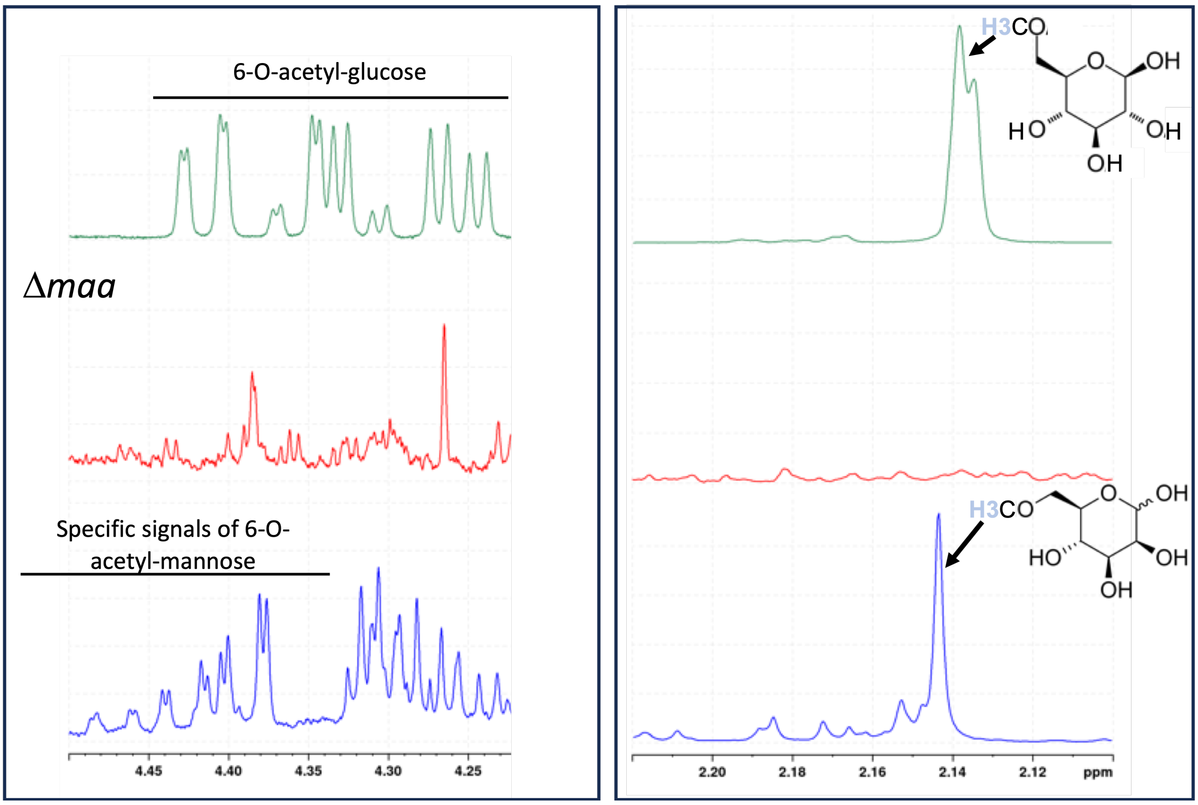


Figure S4. Selected regions of the 1D $^{1}$H NMR spectra of culture supernatants from PFKA1-pBAD-Teth514_1788 (blue) and PFMA1-pBAD-Teth514_1788 (red), compared with spectra of a 6-O-acetyl-glucose standard (green). As 6-O-acetyl-mannose is not commercially available, 6-O-acetyl-glucose was used as the closest structural analogue for metabolite identification. Maltose O-acetyltransferase (encoded by *maa*) has been shown to acetylate a broad range of sugars, including mannose (Brand and Boos, 1991). The absence of this metabolite in the PFMA1 strain, resulting from maa deletion, supports its tentative identification as 6-O-acetyl-mannose.

Brand B, Boos W. Maltose transacetylase of Escherichia coli. Mapping and cloning of its structural, gene, mac, and characterization of the enzyme as a dimer of identical polypeptides with a molecular weight of 20,000. J Biol Chem. 1991 Jul 25;266(21):14113-8. PMID: 1856235.

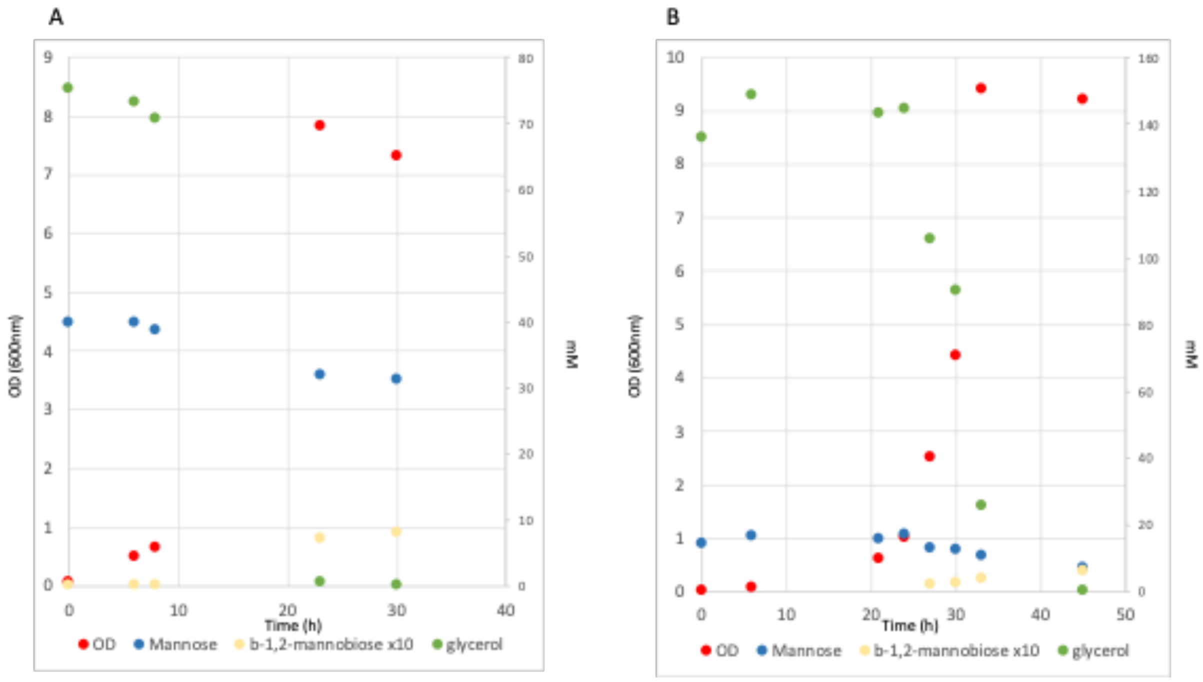


Figure S5. Growth profiles of *E. coli* PFMA1-Teth514_1788 strains cultivated in bioreactors on mannose-glycerol mixture. A. Mannose 7 g.L$^{-1}$; glycerol 7 g.L$^{-1}$. B. Mannose 2.5 g.L$^{-1}$; glycerol 12 g.L$^{-1}$.
